# Sequencing of clinical samples reveals that adaptation keeps establishing during H7N9 virus infection in humans

**DOI:** 10.1101/2020.12.30.424890

**Authors:** Liqiang Li, Jinmin Ma, Jiandong Li, Jianying Yuan, Wei Su, Tao jin, Xinfa Wang, Renli Zhang, Rongrong Zou, Lei Li, Jianming Li, Shisong Fang, Jing Yuan, Chentao Yang, Yanwei Qi, Qi Gao, Jingkai Ji, Kailong Ma, Guangyi Fan, Na pei, Yong Deng, Yang Zhou, Dechun Lin, Fei Li, Wenjie Ouyang, Huijue Jia, Xin Liu, Hui Jiang, Huanming Yang, Xun Xu, Hui Wang, Yingxia Liu

## Abstract

The H7 subtype avian influenza viruses (AIV) have a much longer history and their adaptation through evolution pose continuous threat to humans ^1^. Since 2013 March, the novel reasserted H7N9 subtype have transmitted to humans through their repeated assertion in the poultry market. Through repeated transmission, H7N9 gradually became the second AIV subtype posing greater public health risk after H5N1 ^2,3^. After infection, how the virus tunes its genome to adapt and evolve in humans remains unknown. Through direct amplification of H7N9 and high throughput (HT) sequencing of full genomes from the swabs and lower respiratory tract samples collected from infected patients in Shenzhen, China, we have analyzed the *in vivo* H7N9 mutations at the level of whole genomes and have compared with the genomes derived by *in vitro* cultures. These comparisons and frequency analysis against the H7N9 genomes in the public database, 40 amino acids were identified that play potential roles in virus adaptation during H7N9 infection in humans. Various synonymous mutations were also identified that might be crucial to H7N9 adaptation in humans. The mechanism of these mutations occurred in a single infection are discussed in this study.

---

Earlier study have shown that after just one or two normal passages propagation in chick embryos, human Influenza virus A differs sharply from the *in vivo* ones ^4^. Not only were tissue tropism and binding properties altered, but the mutations responsible also gave rise to detectable changes to their antigenic characteristics. Influenza A virus culture-based mutation phenomenon were also reported by several studies ^5-8^. Since the first outbreak of H7N9, full genomes of various isolates derived human have been sequenced. Genomes of most of the human H7N9 viruses were obtained after *in vitro* propagation in embryonated chicken eggs. Ren *et al*. also found that the consensus genomes from the original human patients and the embryonated eggs culture show differences^12^. Previous studies have shown that H7N9 viruses show mutations during infecting the ferret ^9-11^. Over time, ranging from 12.6-40 days ^13^, H7N9 virus in infected human host should have tuned their genetic information for adapting to novel mammalian host. We hypothesize that characterizing the genomes of H7N9 viruses *in vivo*, other than in vitro cultures can help to find out the mutations that might be playing roles during adaption in the host, and observing the intra-host characteristics.

To verified this thought, we analyzed samples derived from H7N9 infected patients admitted to the Third People’s Hospital of Shenzhen (TPH-SZ). Nasal and pharyngeal swabs and phlegm samples from four patients (one mild and three severe patients) were collected and subjected to HT sequencing with and without embryonated chicken egg propagation (Supplementary Table 1, 2; Supplementary discussion) (Phase I). Surprisingly, pairwise alignment of the above 4 pair H7N9 isolates showed that consensus sequences changed for the same sample before and after *in vitro* culture, with identities varying from 0% to 4.8% on each segment (Figure 1a). The polymerase encoding segments, segment 1-3, maintained high identity following culture, when compared to the other five segments (5-8) (Figure 1a; supplementary table 3). The genetic variation in the NP and NS was the largest, showing rapid diversification, consistent with the previous findings in human at macro-evolution level ^14^. One pair (2014S4/cul.2014S4) maintained all 8 consensus genome segments well, with the exception of PA and M segments, which showed changing only 0.1% and 0.6% variation, respectively (Figure 1a; supplementary table 3). The phylogenetic relationship based on the later five genome segments were also changed when compared against the isolated derived from *in vitro* egg culture (Supplementary figure 1). These sharply changes from *in vitro* propagation were also verified by compared our data in this study with cultured Shenzhen human H7N9 and avian H7N9 isolates report in Lam *et al*. (Supplementary discussion).

**Figure 1.**
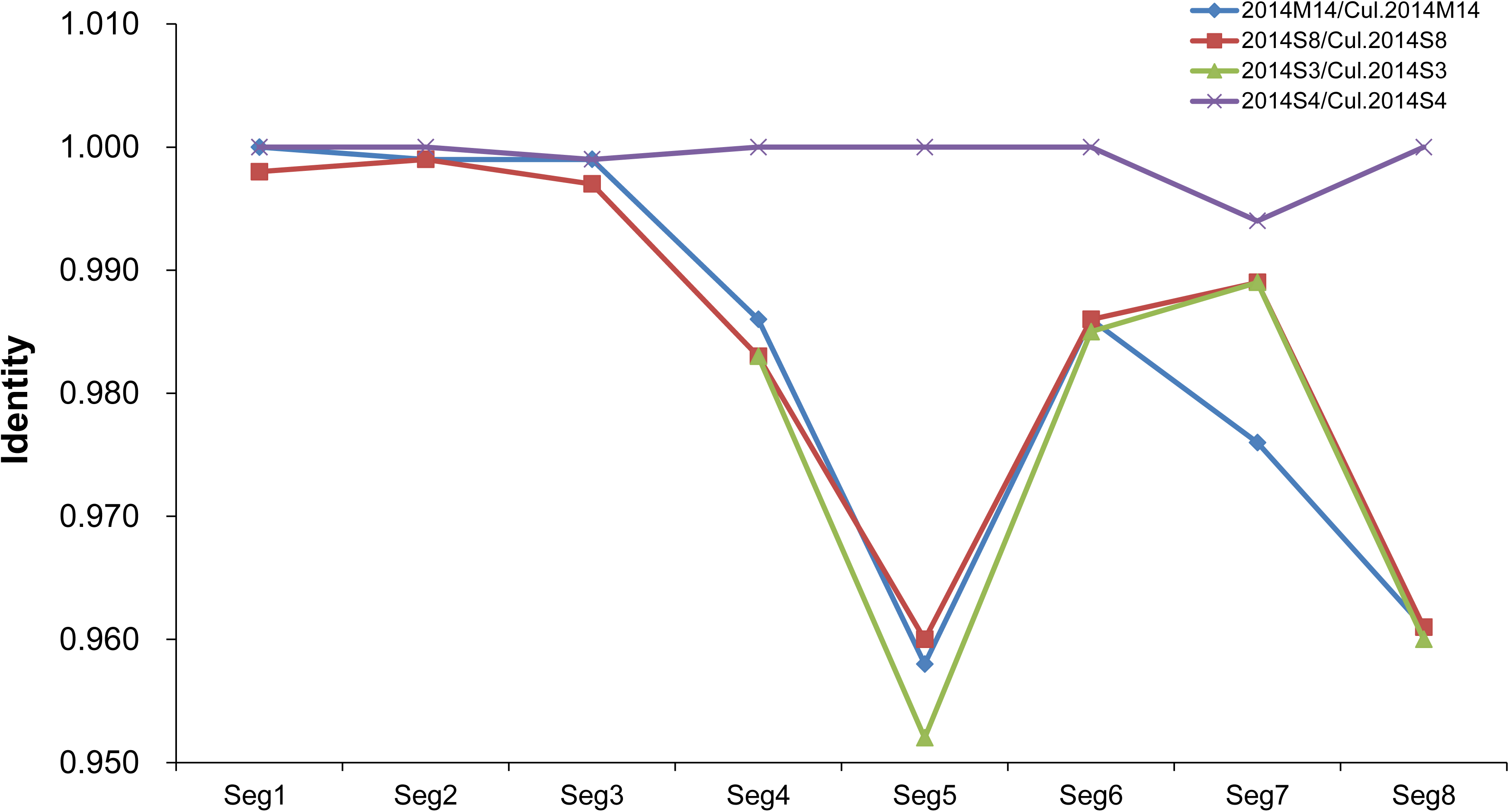
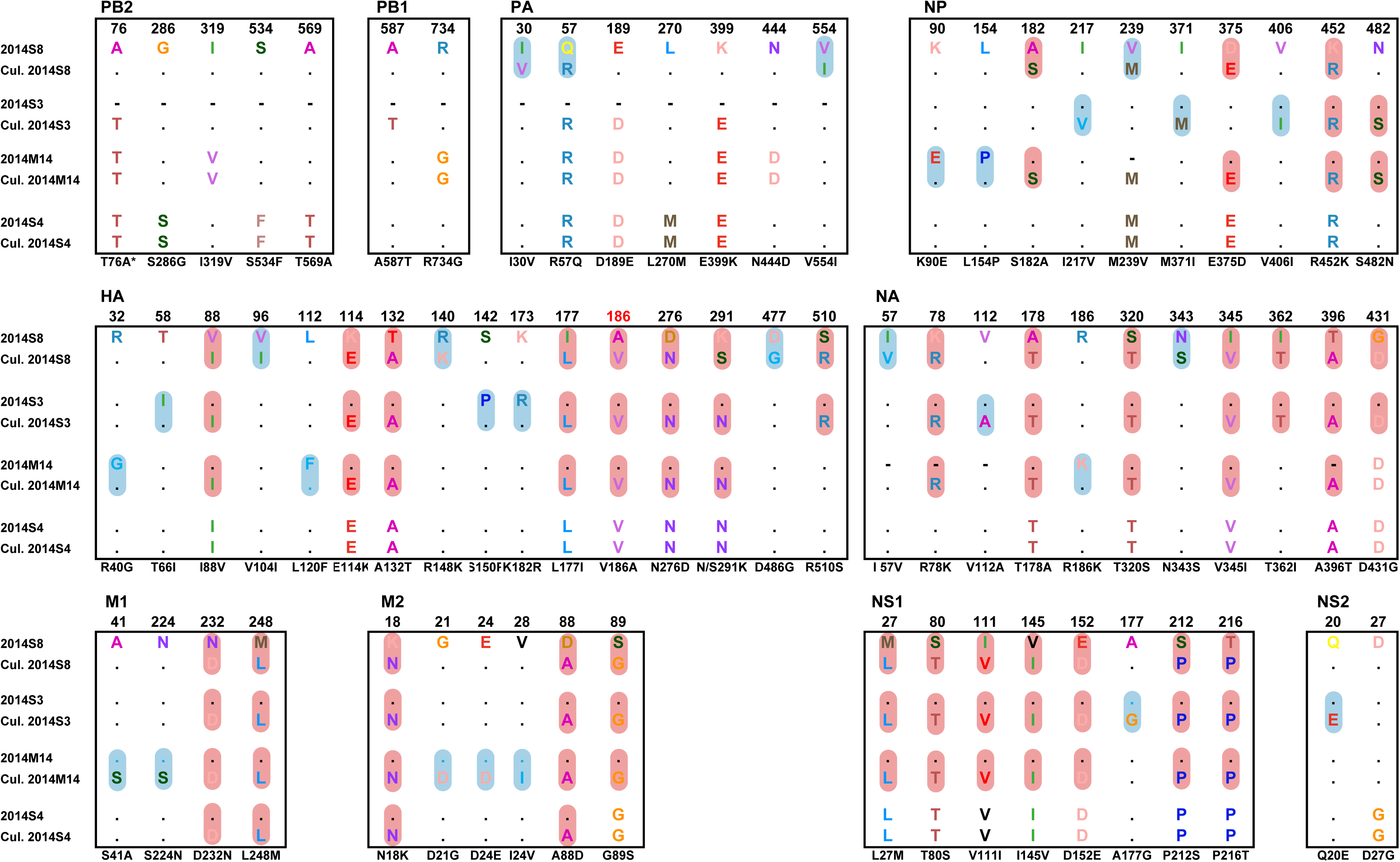
Comparison of the human H7N9 consensus sequence obtained from direct clinical samples and embryonated chicken egg cultures in phase I. a), Identity of consensus nucleotide sequences between four *in vivo* H7N9 viruses and their *in vitro* cultured counterparts; 8 genomic segments are list in x-axis sequentially, y-axis are nucleotide sequence identities of each paired samples. b) Amino acids mutations identified by the four cultured and uncultured pair human H7N9 virus samples. Residues mutations occurred in more than two pairs are shaded in pink, singleton mutations are shaded in blue. *, T76A, the residue T, before the number 76 is putative specific in avian host, and the residue A after 76 is putative specific in avian host.

For the three high-mutated pairs, on average about 1 out of 4 mutations were non-synonymous, 3 out of 4 were synonymous (Supplementary table 4). Surprisingly, most of these mutations were convergent (appeared at least in two isolates) (Supplementary table 4). Through depth frequency analysis, we found that most of these mutations were largely dominant (depth frequency >99%, supplementary table 6,7), suggesting that these mutations can be established following just one passage (or once infection in human). From neutral evolution theory and by comparing with egg-cultured isolates in parallel, we found that the H7N9 NP, NS1/2, M1, and HA2 suffered the strong purification effect, while the polymerase complex components PB2, PB1, PA, and the two surface proteins HA1 and M2 are in a steady neutral evolution during host shift (supplementary table 4,5).

Further, non-synonymous mutations, e.g. amino-acid level changes were observed in 75 residues compared with the egg cultures ones. 33 of them (41.3%) showed convergent (Figure 1b). We postulate that these 33 residues might be more adaptable in humans and play a vital role during establishment and evolution of infection in humans. To verify this, additional 35 isolates were further analyzed by HT sequencing from the H7N9-infecting patients collected from the 2014 and 2015 pandemic season (Phase II). Data of 18 isolates directly from patients, 6 isolates of egg cultured human H7N9 (the 2013-2014 season) were obtained (Supplementary table 2). Overall, all 33 residues occurred in human *in vivo* H7N9 viruses (SZ_*in-vivo*-H7N9, n=22) at a higher frequency (>30%) than in cultured human H7N9 viruses (SZ_*in-vitro*-H7N9, n=10), with 28 residues showing statistics significance (Figure 2a). Between 2013-2014 and 2014-2015 endemic seasons (11 vs 11) of SZ_*in-vivo*-H7N9, occurring frequencies showed no statistics differences except at 89S (M2) but all significant higher than SZ_*in-vitro*-H7N9, ruling out the influence of genetic background (Figure 2a). We also compared the occurrence frequency of these 33 shift residues between these SZ_*in-vivo*-H7N9 with the avian and human H7N9 sequences deposited in the GenBank (GB_avian-H7N9 and GB_human-H7N9) since 2013 in China. 22 residues in SZ_*in-vivo* H7N9 showed higher occurrence frequencies than in GB_avian-H7N9 (n=∼400), and 19, included in the above 22 than GB_human-H7N9 (n=∼80), and 21, included in the above 22 than SZ_*in-vitro*-H7N9 with statistical significance (Figure 2b). Unexpectedly, most (22∼23) of these 33 residues occurring frequencies showed no significant differences among the GB_avian-H7N9, GB_human-H7N9 and SZ_*in-vitro*-H7N9, except the residues in NP and NS1 segments (Figure 2b).

**Figure 2.**
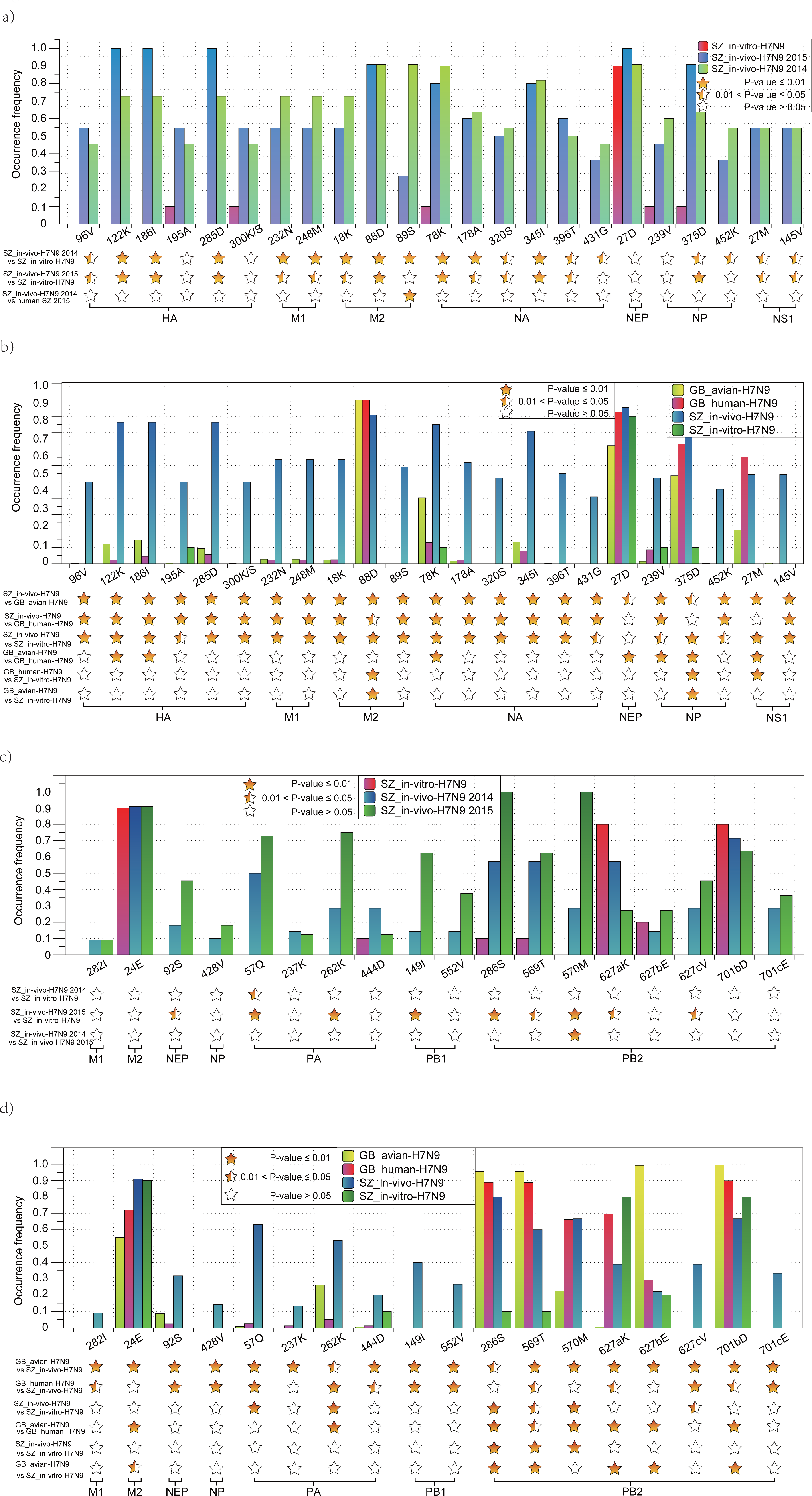
Occurring frequency comparison of mutation residues among SZ_*in vivo-*H7N9, SZ_*in-vitro*-H7N9, and GB_avian-H7N9 and GB_human-H7N9 deposit in GenBank. a), comparison of convergent mutation residues (phase I) between SZ_*in-vivo*-H7N9 (n=∼22), SZ_*in-vitro* H7N9 (n=10); b), comparison of convergent mutation residues among SZ_*in-vivo*-H7N9, SZ_*in-vitro*-H7N9, and GB_avian-H7N9 (n=∼99-411) and GB_human-H7N9 (n=∼80-100); c), comparison of residues showing population bias between SZ_*in-vivo*-H7N9 and SZ_*in-vitro*-H7N9; d), comparison of residues showing population bias (from phase II) among SZ_*in-vivo*-H7N9, SZ_*in-vitro*-H7N9, GB_avian-H7N9 and GB_human-H7N9. Y-axis is occurring frequency (percentage) of mutation residues. Occurring frequency differences among different groups were compared using fisher exact test. Alignment files can be provided upon request.

Besides the above convergent mutations in four paired samples, we further identified other variable sites between SZ_*in-vivo*-H7N9 and SZ_*in-vitro*-H7N9, and then compared the mutations of these sites with GB_avian-H7N9 and GB_human-H7N9. Among the 55 variable residue sites, though only 1 site (57Q of PA) in SZ_*in-vivo*-H7N9 showed significant higher frequency than SZ_*in-vitro*-H7N9 (Figure 2c), mount to 18 of the 55 residue sites showed higher occurrence frequencies than in GB_avian-H7N9 and 13 of them were higher than GB_human-H7N9 (Figure 2d). Simliar to the convergent mutations identified in phase I, most (9-12) of 13 residues showed no differences among GB_avian-H7N9 and GB_human-H7N9 and SZ_*in-vitro*-H7N9. Because most viruses of GB_human-H7N9 were derived through *in vitro* cultures, we postulate that human H7N9 have reversed to avian H7N9 status after *in vitro* culture, a trait not explained before. This sequence reversal could be attributed to the viral adaptation in the embryonated eggs.

Collectively, the above results reveal that the *in vitro* culture has considerable influence on the H7N9 at the genome, viral proteins. 40 residue sites (Supplementary table 8) were identified in a high frequency (27-100%, Figure 2) in human *in vivo* H7N9 virus. All these mutations were also verified by Illumnia Hiseq 2500 sequencing (data not shown). Most of these mutations are not identified playing certain function in previous studies. Primary structure mapping and tetra-structure remodeling analysis were done to assess the functional influences of these mutations. Two surface glycoproteins, HA and NA contain the most mutations, amount to 7 on HA, and 6 on NA. In HA, all mutations are located in HA1: 3 (114K, 177I and 186A) were located near the receptor-binding pocket (Figure 3), 2 (88V, 273I) in the esterase subdomain,and 2 (276D, 300K) in the fusion domain. Due to their functional locations, these mutations could play a significant biological role in HA stability, cell fusion and/or receptor-binding specificity or affinity. In NA, all 6 (78K, 178A, 320S, 345I, 396T, 431G) mutations were located on the head domain and close to the C-terminal, with 78K located near the stalk and trans-membrane region. These mutations might play important roles in HA-NA interaction and virion stability.

**Figure 3.**
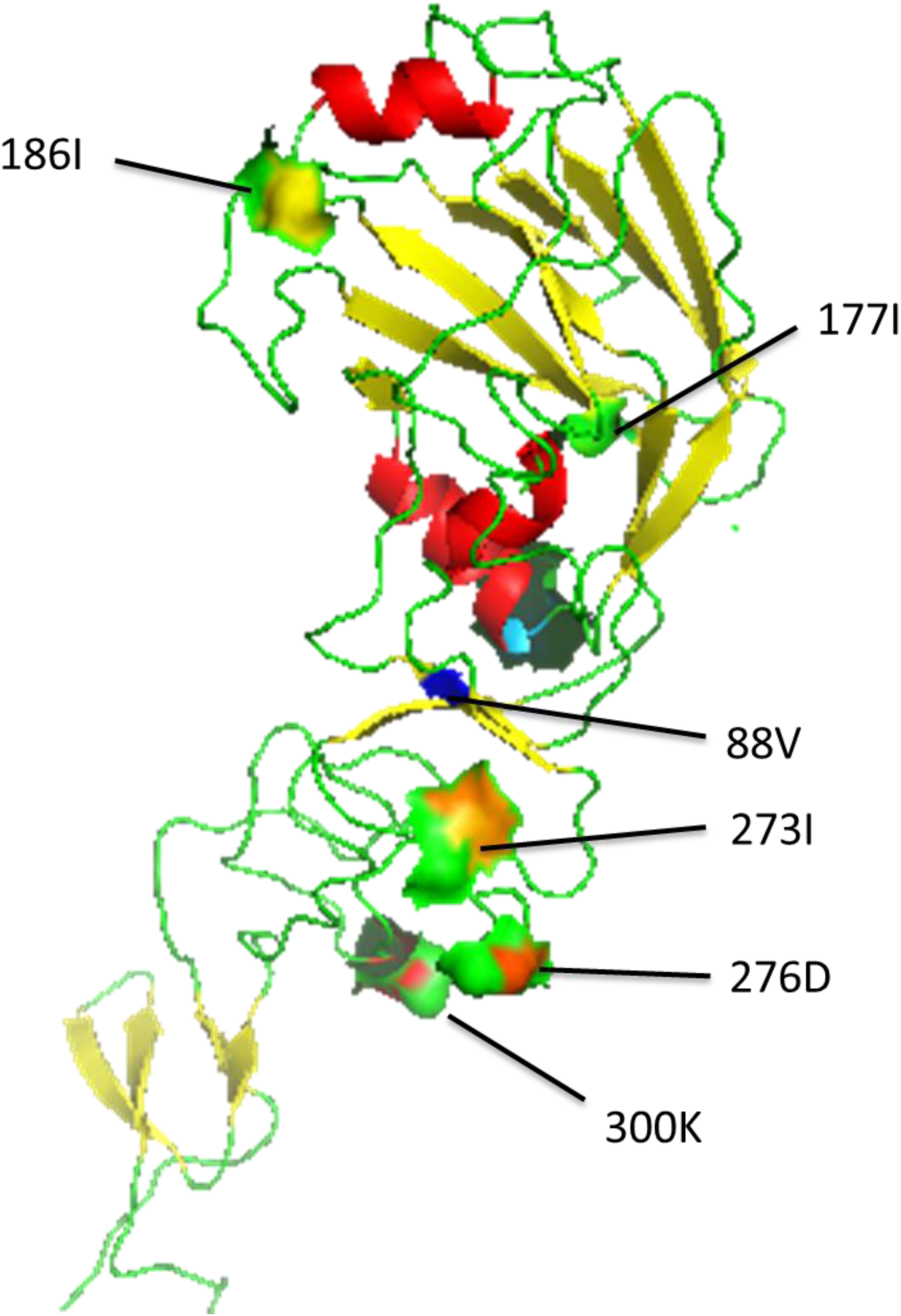
Structure modeling of mutation sites in HA of human H7N9. Structural modeling were generated using the structural files of two mature proteins, HA1 (PDB ID 4ln6.1.A) and HA2 (4bsa.1.B) in PyMOL viewer.

When AIV jumps to a new host, sustainable and stable replication is its first key steps to survival, adaptation and establishment of infection ^15^. For the virion replication complex, ribonucleoproteins (RNP), mutations mostly occurred at the domains involved in RNP protein interaction (Figure 2; supplementary figure 3), might playing an important role in effective and stable replication establishment in human. It is know that PB2 plays key roles during host adaption. 570M mutation was identified located in the CAP binding domain and near the C-terminal NP and PB1 interaction domain. Especially, at residue 627 and 701, besides the well-known host adaption mutation 627K, 701N, two novel mutations, 627V and 701E, were also identified occurring at high frequency in *in-vivo* H7N9 respectively (Figure 2d), suggesting might be adapted in human host. In PB1, 2 mutations were identified, one located near the N terminal and the other in the PB2 interaction domain near the C-terminus (supplementary figure 3). As for the PA, 4 mutations were identified, 57Q localized at the N terminal, two in (237K)/near (262K) the NLS domain, and 444D at the middle of the PA. As for the NP, 4 mutations were identified, all localized in the NP-NP interaction domain, with 3 of them belonging to the NP-PB2 interaction domain implying roles in NP-NP complex stability in virus replication (supplementary figure 3).

It is believed that the viral matrix protein M1 plays an important role during RNP nuclear transportation ^16^, NS1 is the main viral antagonist of the innate immune response during influenza virus infection to overcome the first barrier the host presents to halt the viral infection ^17^. 5 mutational changes were identified in the matrix protein, 2 (232N, 248M) were located at the RNP interaction domain on M1, 2 (18K and 24E) were closely located at the N terminal of M2 with 24E belonging to the RNP interaction domain, and 1 (89S) was located close to the N terminal of M2 (Supplementary figure 3). The distributions of these mutations are consistent with the possible role they may have in viral RNP complex transport, viral packaging and budding. 3 mutations were identified in the NS1/NEP, 2 of which resided in the NS1 with 27M resident at N-terminal dsRNA/PABP1/RIGI/EIB-AP5 binding domain and 145V in the short intergenic region of CPSF30 and NES domain, the third mutation (92S) located in C-terminal of NEP (Supplementary figure 3). These mutations might play roles in RNP transportation or in escaping the host immune response.

In recent years, synonymous mutations, correlating with the codon usage bias issues, were believed to influence virus fitness and pathogenicity through translation efficiency regulation and CpG and TpA dinucleotides frequency changes ^18^. G+C content, CpG and ApT dinucleotide frequency, GC3 number and observed ENcs changed during host shift (supplementary figure 5). Through phase I four paired sample comparison, we found that, unlike the non-synonymous mutations in which A-C transversions (tv) occurred at a much higher frequency besides purine (A/G) to purine transitions (ts), the synonymous mutations showed that that (A/G), or pyrimidine (C/T) to pyrimidine tv occurred at a frequency higher than the purine to pyrimidine or pyrimidine to purine tv similar to other organisms ^19^(Supplementary figure 4). Further, most of the synonymous mutations are convergent across different pairs and not distributed randomly and uniformly on the genome segments, especially one to four synonymous mutations cluster found on the HA, NA, M1 and NS1 segments (Supplementary figure 9). Collectively, these changes suggest H7N9 virus might be tune their genome to adapt to human, and might play important roles in viral replication and immune escape during H7N9 infecting human.

To analyze how these viruses has been influenced by these accompanying mutations, we compared their intra-host diversity and show that *in vitro* H7N9 viruses shows less diversity than *in vivo* human H7N9 viruses suggesting bottle-neck effect to have occurred during *in vitro* culture (Supplementary figure 7a), and display a different diversity profile (Supplementary figure 6). Most human adapted mutations might have been filtered during inoculation to the egg, the second host jump, and the minority egg suitable ones quickly arise during *in vitro* culture. Intra-host diversity correlation analysis showed that in human the NP and NS1/2, MP segments are closely linked, otherwise in embryo chicken, the HA are closely related with MP and NS1/2 (Supplementary figure 8), suggesting different intra-host fluctuation mechanisms. Thus, H7N9 might have utilized the unknown molecular strategy for adaptation, efficient replication and establishment in a new host.

In summary, we found that *in vivo* H7N9 virus in the human host are enduring a genetic tune to adaption and in this context, we, identified various amino acids and synonymous mutations that might play a critical role in infection. This gives us a different aspect to recognize AIV infecting human or other mammal hosts and provide vital clues for further experimental and functional validation *in vitro*. We also found *in vitro* culture can change the intra-host diversity of *in vivo* H7N9 virus and produce various mutations during a single passage. These findings provide further molecular insights into functional identification of virus *in vivo* and design next generation of therapeutic strategies to halt virus infection at critical stages of life cycle in humans.

## Supporting information

Supplementary discussion

## Acknowledgements

We thank Professor Nitin K. Saksena for his constructive work on manuscript revising (Sydney Medical School, the University of Sydney). This study was supported by Shenzhen Science and Technology Research and Development projects (Grant No. JCYJ20150402111430617 and JCYJ20151029151932602), Key specialized fund for new infectious diseases in Shenzhen City (No. 201161).

## Author contributions

The manuscript was written by L.Q.L., H.W, J.M.M, Y.X.L, G.F.G. and W.-C.C. Samples were collected by R.R.Z, T.J, R.L.Z, Y.M.L, S.S.F, X.F.W. Experiment and data analysis were performed by L.Q.L., J.D.L., J.Y.Y., W.S, J.M.M, H.W, C.T.Y, Y.W.Q, Y.X.L, X.L., H.J.J. The study was designed by Y.X.L, L.Q.L, H.W, X.X.

## Methods

### Patients and Sample collection

From Dec 18th, 2013 to April 1st, 2014, 25 H7N9 infection patients with Influenza-like symptoms in Shenzhen City, Guangdong Province were admitted to the Third People’s Hospital of Shenzhen (TPH-SZ). In the second waves of H7N9 outbreak in Shenzhen, i.e. third wave in China, 13 more patients infected with H7N9 were confirm in Shenzhen and were admitted to TPH-SZ from Jan. 4th, 2015 to April 2nd, 2015. In all these cases, diagnostic and treatment decisions were made by consortia of more than three panel members of clinical specialists. After admitted, we immediately collected the respiratory sample for clinical confirmation of H7N9 infecting. Standard Real-time reverse transcription polymerase chain reaction (RT-PCR) assay for H7N9 confirmation were done in the Influenza Reference Laboratory of the Shenzhen municipal center for disease control (CDC) and Guangdong province key laboratory of emerging infectious diseases., C_T_ value smaller than 38 were judged as H7N9 positive, according to the Guidelines authorized by China National Influenza Center of China CDC. Mild and severe cases were distinguished according to IDSA/ATS criteria.^1^ Severe cases met at least one of major criteria (Invasive mechanical ventilation/Septic shock with the need for vasopressors) or more than three minor criteria ^1^. Analysis of the patient’s clinical samples for the identification of potential pathogens was approved by the ethics committee of TPH-SZ.

### Viral RNA preparation and high throughout Sequencing

Nasopharyngeal swab, phlegm samples and some other respiratory specimens were collected for diagnosis and egg culture. Nasal, oropharyngeal swabs and/or tracheal aspirate samples from each patient were collected into transport medium and separated into two aliquots within two hours, one for diagnostic labs and the other for long term storage. For in vivo analysis, viral RNAs were extracted directly from pharyngeal and nasal swabs or phlegm samples using Qiagen viral RNA mini kit. For *in vitro* virus isolation, samples testing positive for both the H7 and N9 genes were propagated in 10-day-old embryonated chicken eggs and then cultured at 37 °C for 2 days in a bio-safety level 3 laboratory. Viral RNA were extracted using Qiagen viral RNA mini kit from allantoic fluid, and tested for H7 and N9 genes by RT-PCR assays and, if positive, used for further genomic sequencing.

To obtain whole H7N9 genome, reserves transcriptions were done from viral RNA samples using primer pair uniR_RT (5’-AGTAGAAACAAGG-3’) and uniF_RT (AGCGAAAGCAGG) to obtain viral full genomic cDNAs. Then PCR application using specific primer pairs (Supplemental Table 2) designed for H7N9 virus and Takara One-step RT-PCR kit (Takara, China) segment by segment. PCR products of 8 segments were quantified by 1% gel, and mixed with equal mole and then subject to library construction and HT sequencing using BGI seq100 (Ion proton, Life technology, USA).

### Consensus genome construction, comparison and phylogenetic analysis

To obtain the consensus genome, we mapping all reads to a reference H7N9 virus with the threshold match rate at 0.80, and at each site, we choose the dominant bases as final consensus genome. Sequences were aligned using MUSCLE v3.5. Phylogenies were inferred on the basis of Neighbor-Joining method, by using the maximum composite likelihood model in MEGA 6.0 ^2,3^. Topological robustness was assessed by bootstrap test. For analysis the novel mutations in human H7N9, we retrieved all protein sequences of avian and human H7N9 genome segments with full length ORF from GenBank (until 08 Sep. 2015).

### Entropy, mutation rate, amino acid mutation rate and viral fitness

Intra-host diversity implication viral fitness were assessed by the average Entropy or mutation rate of one genome segment or whole genome. Entropy were calculated by using equation, S = −100 * Sum (P*i* * log P*i*) where P*i* is the frequency of the *i*th allele ^5^. Rate of minor nucleotide variant allele, other than the base with the largest depth, at each nucleic acid site were calculated by the minor allele depth deduced the total depth of this site. Mutation rate of each nucleic acid site were designate the ratio of the minor variant allele, that is adding all Rate of all Minor variant allele (MR*i*). Minor alleles changing the amino acid coding were designated as a no-synonymous SNV. At specific codon, the sum value of each no-synonymous nucleotide mutations rate compared to consensus codon were designated as the at one amino acid site were add together were designated as amino acid mutation rate.

Codon-based test of neutrality were done using Nei-Gojobori method in MEGA 6 ^3^. G+C content, CpG and ApT dinucleotide frequency, GC3 number and observed ENcs were calculated using SSE version 1.2 ^4^.

### Linear mapping and tertiary structure of amino acid mutations

The linear primary structural maps of HA, NA, PB2, PB1, PA, NP, M1/2 and NS1/NEP were derived and modified from the previous maps of Ping, J., et al.^5^. Tertiary structures were generated using the PyMOL viewer. For H7, Structural maps were generated using the structural files of two mature proteins, HA1 (PDB ID 4ln6.1.A) and HA2 (PDB ID 4bsa.1.B).

## Supplementary material legend

### Supplementary figures

**Supplementary figure 1. Phylogenies relationship of hemagglutinin** (a), **neuraminidase and PB2** (b) **genes of 4 H7N9 clinical-culture pairs**. Bootstrap support values (%) from 1,000 replicates are shown for selected lineages. The scale bar to the left of each tree represents the substitutions per site. Four phase I in-vivo and in vitro H7N9 pairs were indicated by colored solid circle.

**Supplementary figure 2. Sequence Comparison of SZ_*in-vivo*-H7N9 and SZ_*in-vitro*-H7N9 with other Shenzhen human and avian H7N9 virus reported in Lam *et al***. a), heatmap and cluster analysis of pairwise identity data; b), with-in genetic diversity and between-group distance analysis of SZ_*in-vivo*-H7N9 and SZ_*in-vitro*-H7N9 with other Shenzhen human and avian H7N9 virus reported in Lam *et al*.

**Supplementary figure 3. Linear mapping of potential functional adaptive sites to primary structure of H7N9 structural and functional proteins**. Boxes represent viral proteins are not drawn according to the actual length ratio.

**Supplementary figure. 4. Comparison of transition and transversion mutation frequency of four *in vivo* H7N9 and their *in vitro* cultured counterparts**.

**Supplementary figure 5. Codon usage differences of four paired *in vivo* H7N9 compared with *in vitro* H7N9 virus**. Cells indicated with red, green and grey colors represent up, down and no significant change respectively, compared with *in vitro* H7N9 virus (fisher exact test).

**Supplementary figure 6. Whole genome intra-host diversity profile of Shenzhen H7N9 influenza viruses**. Heatmap showing nucleotide mutation frequencies of genomic segment identified in SZ_*in-vivo*-H7N9 and SZ_*in-vitro*-H7N9. At each site, bases other than the base with the largest depth were designated as minor alleles. Nucleotide mutation frequencies were calculated by the minor allele depth deduced the total depth. 8 genomic segments are ordered sequentially, corresponding to PB2, PB1, PA, HA, NA, NP, M1/2, NS1/2 respectively.

**Supplementary figure 7. Intra-host diversity of Shenzhen H7N9 influenza viruses**. a, b, Comparison of mean nucleotide mutation frequency of whole genome. Mean nucleotide mutation frequency for each sample was calculated by adding all nucleotide mutation frequencies together then deduced the total length sequenced by Ion proton. Mean nucleotide mutation frequency was expressed as the negative log. Virus isolated were classified into three different patient prognosis groups, SZ_*in-vivo*-H7N9.S groups H7N9 represented patients manifesting severe symptoms but cured finally; and SZ_*in-vivo*-H7N9.M groups represented patients manifesting mild symptoms through the H7N9 infection and recovered; SZ_*in-vivo*-H7N9.D represented patients manifesting severe symptoms and finally dead. \**P*<0.05, two-tailed Mann–Whitney *U-*test. c, d, Comparison of mean nucleotide mutation frequency of whole genome segment by segment.

**Supplementary figure 8. Correlation analysis among different viral genomic segments of *in vivo* and *in vitro* H7N9 viruses**. Pearson correlation coefficient were calculated using average entropy of each genomic segment of every sample from *in vivo* or *in vitro*.

**Supplementary figure 9. Synonymous mutation distributions analysis of four paired *in vivo* and *in vitro* H7N9 viruses**. On each separate figure, above slide windows analysis shows mutation distribution density, low panel shows each mutation concretely.

### Supplementary tables

**Supplementary table 1**. Sample information of patients investigated in this study.

**Supplementary table 2**. High throughput sequencing profiles of H7N9 viruses in this study.

**Supplementary table 3**. Identity of consensus sequence Human H7N9 between *in vivo* samples and from egg cultured ones.

**Supplementary table 4**. Synonymous and non-synonymous mutation statistics of 4 paired H7N9 viruses in phase I.

**Supplementary table 5**. Codon-based test of neutrality for analysis of 4 paired H7N9 viruses in phase I.

**Supplementary table 6**. Base composition of no-synonymous mutated sites in 4 paired H7N9 viruses in phase I.

**Supplementary table 7**. Base composition of synonymous mutated sites in 4 paired H7N9 viruses in phase I.

**Supplementary table 8**. Functional residues list of H7N9 viral proteins identified in this study.

**Supplementary table 9**. Correlation analysis among different viral genomic segments of *in vivo* and *in vitro* H7N9 viruses.

**Supplementary table 10**. Estimates of average evolutionary divergence within and between H7N9 of different hosts.

**Supplementary table 11**. Primers for H7N9 genome amplification in this study.

