## Supplementary discussion for "Sequencing of clinical samples reveals that adaptation keeps establishing during H7N9 virus infection in humans"

**2 Supplementary discussion**

**2.1 Clinical features and treatment of H7N9 patients**

25 patients from Dec 18th, 2013 to April 2nd, 2014, and 13 patients from Jan. 4th, 2015 to April 2nd, 2015 in Shenzhen City of Guangdong Province were diagnosed infection by H7N9 and admitted to TPH-SZ. 36 patients were adults with the age of 31-81 years old, and only 2 patients were 6 years old children. Of the 25 patients, 71.0% (27/38) were severe cases^1^, and 5 patients (8%) failed to recover and died.

**2.2 High throughput sequencing of Shenzhen 2014 and 2015 human H7N9 viruses**

In phase I, nasal and pharyngeal swabs and phlegm samples were collected from 4 patients (one mild patient and 3 severe patients) for direct viral RNA isolation and egg culture (Supplemental Table 1, 2). Viral RNAs from in vivo samples and cultured ones were subjected to whole H7N9 genome application by using H7N9 specific primer pairs (Supplemental Table 2) and ion torrent sequencing. All 4 pair samples were obtained high quality sequencing data with each segment of an average ORF coverage larger than 90%, an average depth larger than 5,000 and N90 depth larger than 1,000, except several internal genes of 2 *in vivo* samples 2014M14, 2014S3 (Supplemental Table 3).

In phase I, additional 35 isolates were further analyzed by HT sequencing from the H7N9-infecting patients collected from the 2014 and 2015 pandemic season (Phase II). Data of 18 isolates directly from patients, 6 isolates of egg cultured human H7N9 (2013-2014 season) were obtained with each segment of an average ORF coverage larger than 90%, an average depth larger than 5,000 and N90 depth larger than 1,000 (Supplementary table 2), like in phase I, except certain internal genes of 7 *in vivo* samples. We found that we can obtain full genomes from cultured samples easily, but hard for in vivo sample especially for the larger internal genes; and it is much easier to obtain from the lower respiratory tract, throat swab or even phlegm than from nasal. We speculate that H7N9 virus are more adaptive in the lower respiratory tract but shedding lower viral loads than avian host, as reported in previous studies.

**2.3 Phylogenetic analysis of *in vivo* H7N9 viruses**

Phylogenetic analysis based on the alignment of all sequenced human H7N9， Shenzhen human *in vivo* and *in vitro* H7N9, and other sequences used in figure 2 of Lam et al.(selected key viruses for illustrating the genesis of H7N9)^2^, showed that all Shenzhen formed a different clade (southern China) belong to from eastern China human H7N9, consistent with previous studies. As expected, in vitro culture has influenced the phylogenetic relationships between human in vivo and in vitro H7N9. Concretely, four paired H7N9 isolates in phase I has been separated pair by pair in the phylogenetic tree (Supplementary figure 1).

**2.4 Comparison of direct sequenced Shenzhen human H7N9 with cultured H7N9 in this study and in Lam et al 2013.**

To further verify the *in vivo* changes of H7N9 genomes, we have compared, segment by segment, our 22 in vivo H7N9, 10 cultured H7N9 with another 17 chicken embryo cultured H7N9 and 20 cultured avian H7N9 isolated in Shenzhen reported in Lam *et al*.^3^. Cluster analysis based on the pairwise identity matrix showed that segments of most *in vivo* H7N9 formed different clades clearly separated from our cultured ones and cultured ones and avian H7N9 of Lam *et al*. though some are scattered in the *in vitro* clades, or *vice versa*, and all cultured and avian H7N9 are close clustered (Supplementary figure 2a).

This separation was also confirmed by between group genetic distance assessments. The distances between in vivo H7N9 and other groups, ranging from 0.0090±0.0016 (NA) to 0.0326±0.0031 (NP), are larger than that among our in vitro cultured H7N9, and *in vitro* cultured H7N9 and avian H7N9 from lam et al, ranging from 0.0029±0.0007 (NS) to 0.0193±0.0019 (NP) (Supplementary figure 2b). We can also found that except the HA and NA genes, *in vivo* H7N9 are more diversity, showing largest average within-group distance, than other groups (Supplementary figure 2b), might suggesting enduring a selection pressure to dramatically tune their genome to adapt to the new host. Both HA and NA are more conserved, showing lower within-group distances and between-group distances than the internal genes, consistent with previous studies^4^.
