## Supplementary figures and images for "Sequencing of clinical samples reveals that adaptation keeps establishing during H7N9 virus infection in humans"

### Supplementary figure 1

Supplementary figure 1

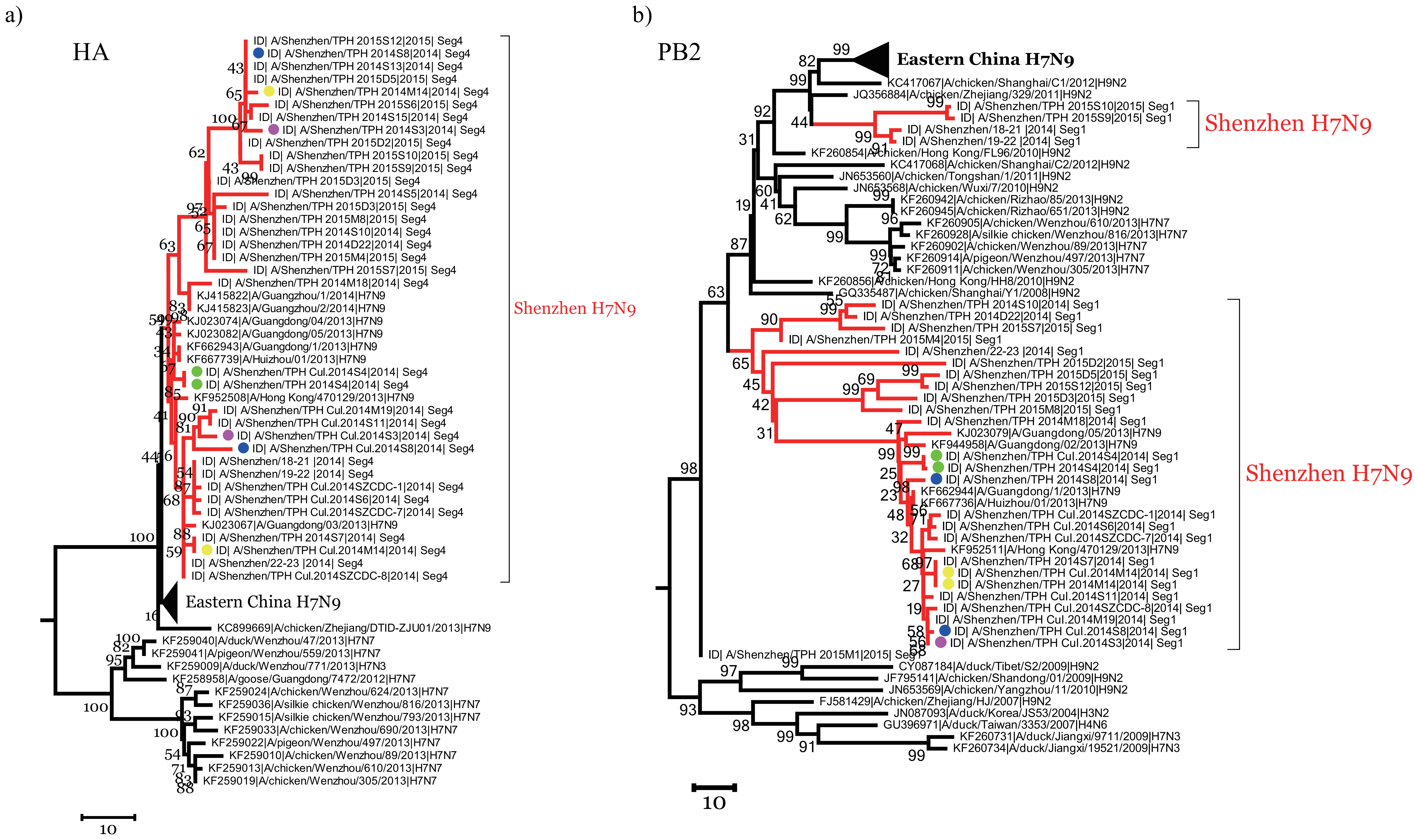

### Supplementary figure 2a

Supplementary Figure 2a

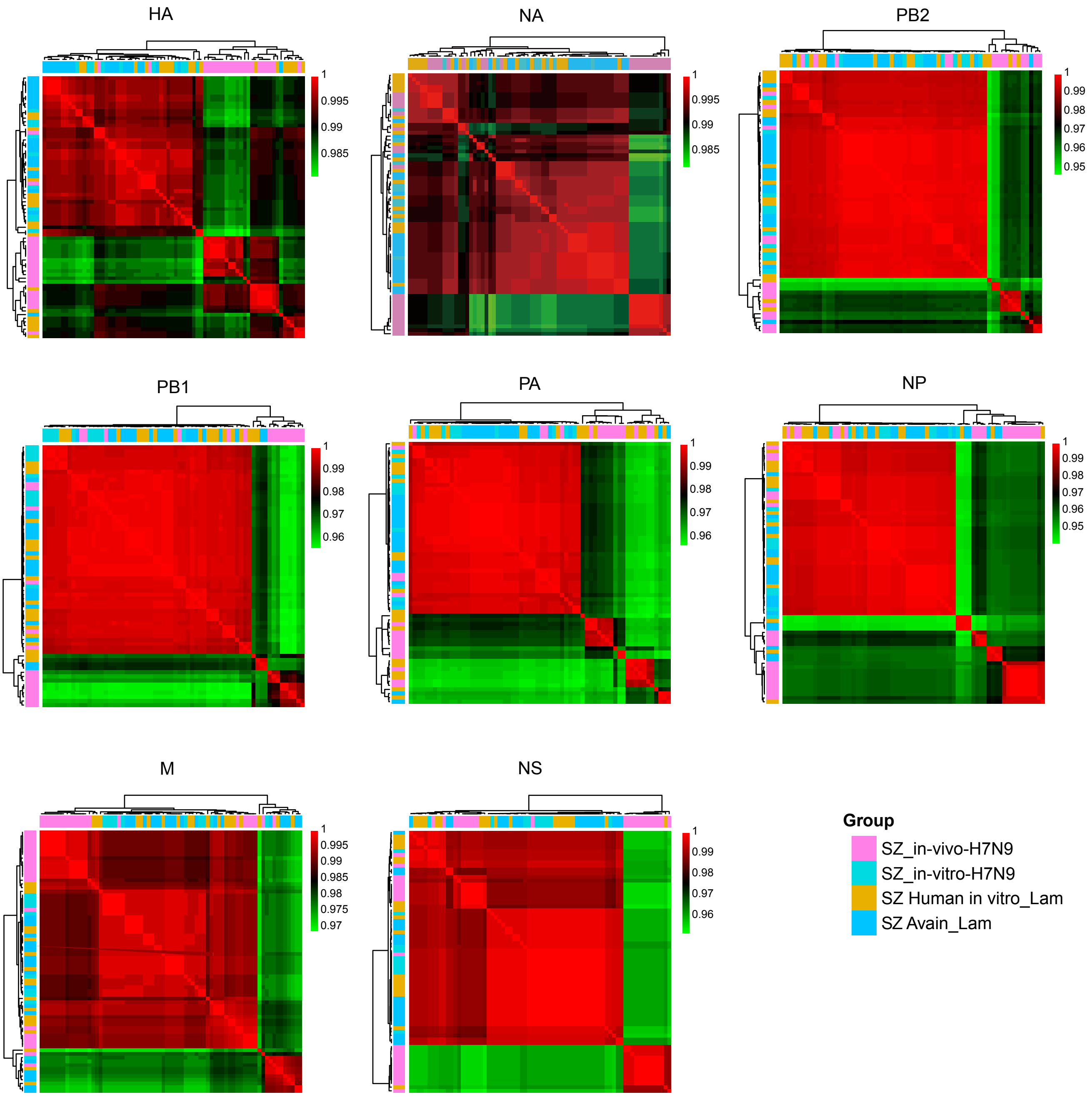

### Supplementary figure 3

Supplementary figure 3

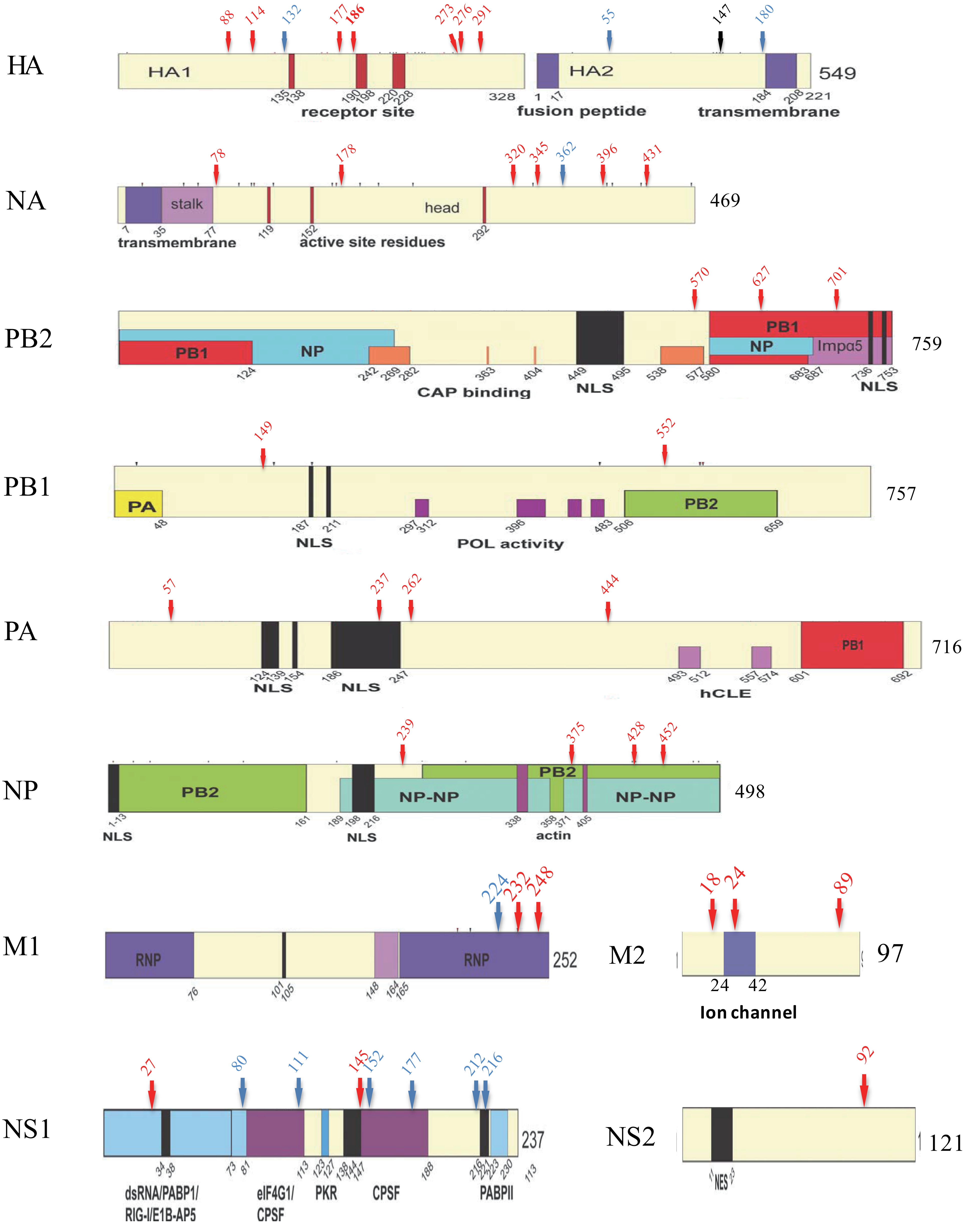

### Supplementary figure 7

Supplementary figure 7

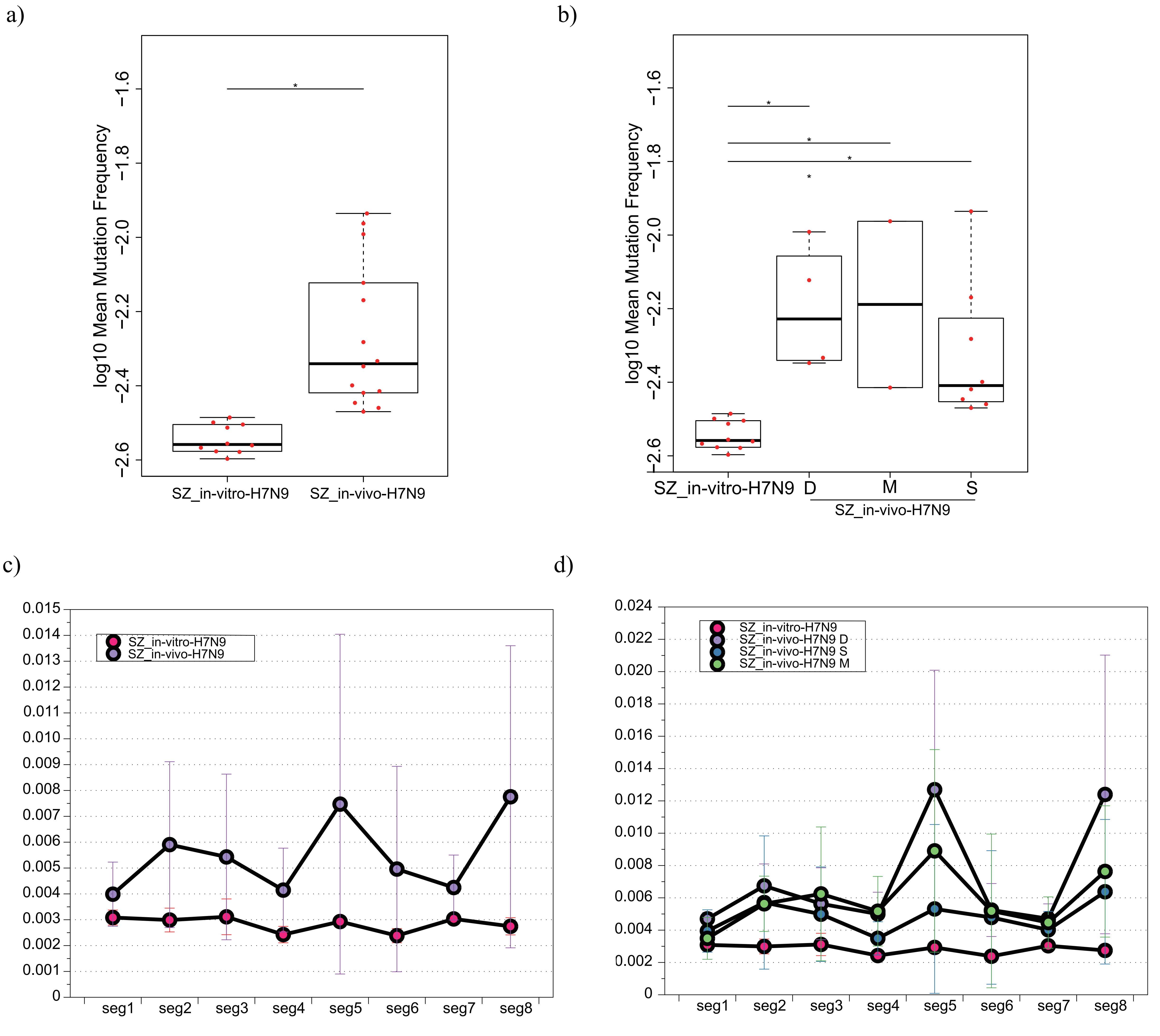

### Supplementary figure 8

Supplementary Figure 8

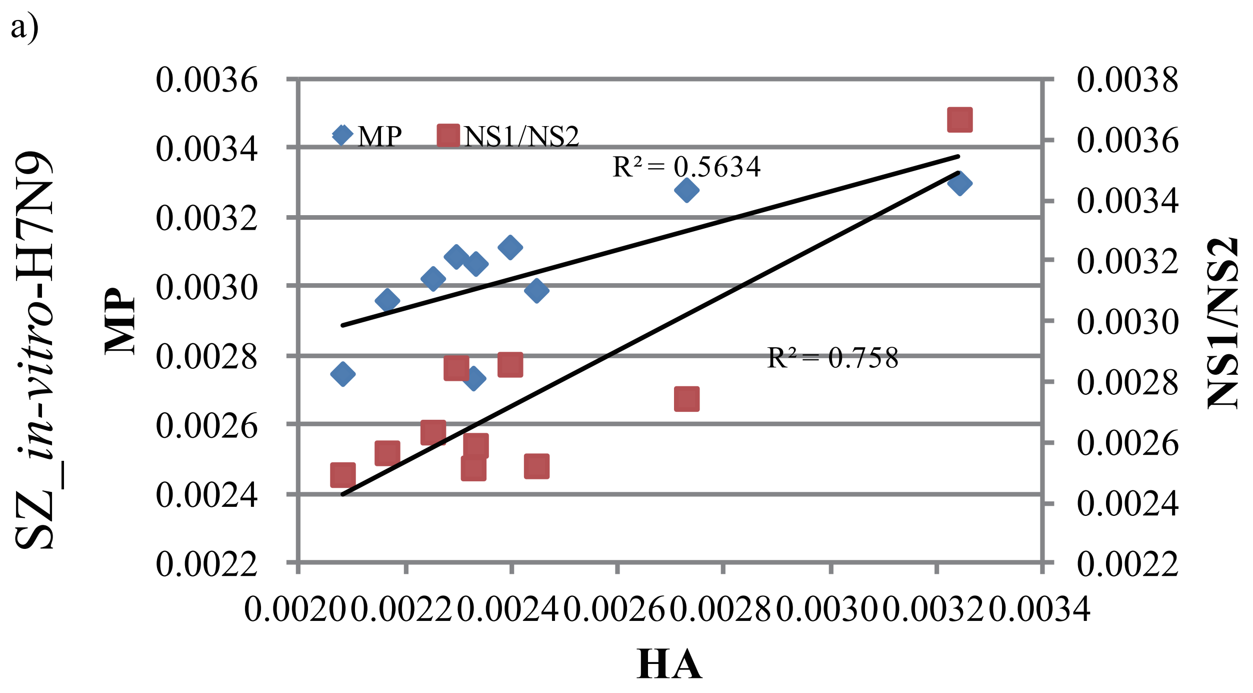


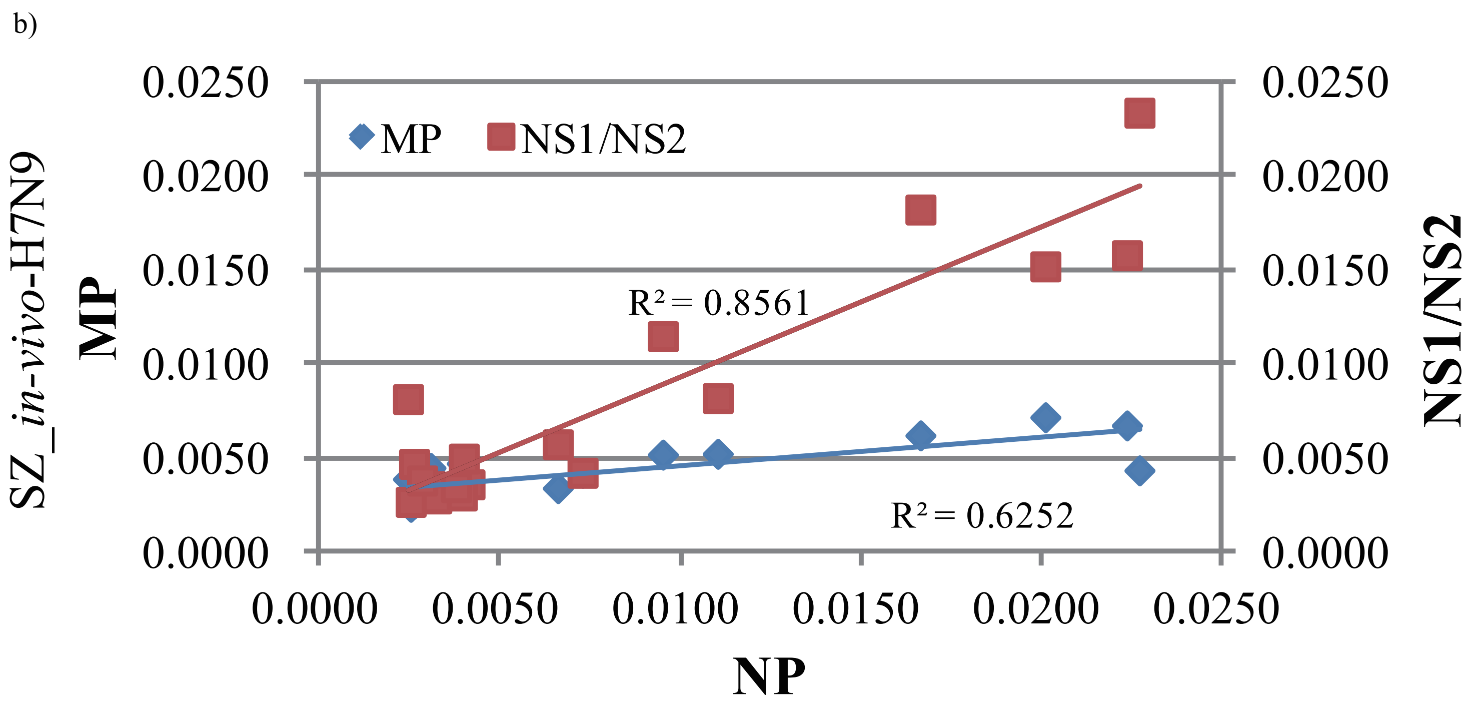

### Supplementary figure 9

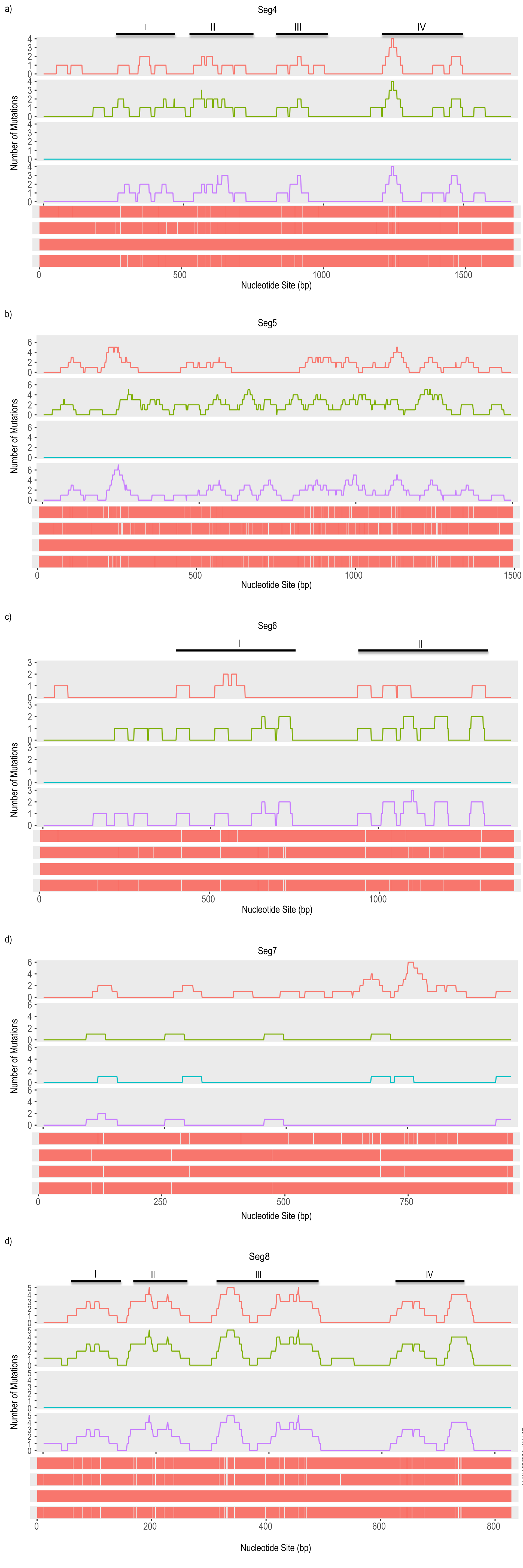
