## Supplementary figure 2b for "Sequencing of clinical samples reveals that adaptation keeps establishing during H7N9 virus infection in humans"

### Within group

| | | Dist | Std. Err |
|---|---|---|---|
| **HA** | | Dist | Std. Err |
| | SZ_in-vivo-H7N9 | 0.0062 | 0.0010 |
| | SZ_in-vitro-H7N9 | 0.0043 | 0.0009 |
| | SZ Human in vitro_Lam | 0.0075 | 0.0011 |
| | SZ Avain_Lam | 0.0034 | 0.0007 |
| **NA** | | Dist | Std. Err |
| | SZ_in-vivo-H7N9 | 0.0070 | 0.0013 |
| | SZ_in-vitro-H7N9 | 0.0039 | 0.0009 |
| | SZ Human in vitro_Lam | 0.0055 | 0.0011 |
| | SZ Avain_Lam | 0.0023 | 0.0006 |
| **PB2** | | Dist | Std. Err |
| | SZ_in-vivo-H7N9 | **0.0312** | 0.0021 |
| | SZ_in-vitro-H7N9 | 0.0027 | 0.0006 |
| | SZ Avain_Lam | 0.0042 | 0.0005 |
| | SZ Human in vitro_Lam | 0.0153 | 0.0012 |
| **PB1** | | Dist | Std. Err |
| | In vivo | **0.0252** | 0.0027 |
| | In vitro | 0.0025 | 0.0007 |
| | SZ Avain_Lam | 0.0062 | 0.0009 |
| | SZ Human in vitro_Lam | 0.0088 | 0.0009 |
| **PB1** | | Dist | Std. Err |
| | SZ_in-vivo-H7N9 | **0.0264** | 0.0024 |
| | SZ_in-vitro-H7N9 | 0.0027 | 0.0006 |
| | SZ Avain_Lam | 0.0080 | 0.0009 |
| | SZ Human in vitro_Lam | 0.0252 | 0.0023 |
| **PA** | | Dist | Std. Err |
| | SZ_in-vivo-H7N9 | **0.0264** | 0.0024 |
| | SZ_in-vitro-H7N9 | 0.0027 | 0.0006 |
| | SZ Avain_Lam | 0.0080 | 0.0009 |
| | SZ Human in vitro_Lam | 0.0252 | 0.0023 |
| **NP** | | Dist | Std. Err |
| | In vivo | **0.0275** | 0.0027 |
| | In vitro | 0.0133 | 0.0015 |
| | SZ Human in vitro_Lam | 0.0169 | 0.0016 |
| | SZ Avain_Lam | 0.0217 | 0.0022 |
| **M** | | Dist | Std. Err |
| | SZ_in-vivo-H7N9 | 0.0075 | 0.0014 |
| | SZ_in-vitro-H7N9 | 0.0053 | 0.0010 |
| | SZ Avain_Lam | 0.0117 | 0.0020 |
| | SZ Human in vitro_Lam | 0.0105 | 0.0014 |
| **NS** | | Dist | Std. Err |
| | SZ_in-vivo-H7N9 | **0.0239** | 0.0039 |
| | SZ_in-vitro-H7N9 | 0.0022 | 0.0007 |
| | SZ Human in vitro_Lam | 0.0091 | 0.0017 |
| | SZ Avain_Lam | 0.0037 | 0.0009 |

### Between group*

| Dist \ Std. Err | SZ_in-vivo-H7N9 | SZ_in-vitro-H7N9 | SZ Human in vitro_Lam | SZ Avain_Lam |
|---|---|---|---|---|
| SZ_in-vivo-H7N9 | | 0.0021 | 0.0016 | 0.0020 |
| SZ_in-vitro-H7N9 | **0.0115** | | 0.0010 | 0.0007 |
| SZ Human in vitro_Lam | **0.0105** | 0.0065 | | 0.0009 |
| SZ Avain_Lam | **0.0110** | 0.0038 | 0.0060 | |
| | SZ_in-vivo-H7N9 | SZ_in-vitro-H7N9 | SZ Human in vitro_Lam | SZ Avain_Lam |
| SZ_in-vivo-H7N9 | | 0.0019 | 0.0016 | 0.0019 |
| SZ_in-vitro-H7N9 | **0.0101** | | 0.0009 | 0.0006 |
| SZ Human in vitro_Lam | **0.0090** | 0.0050 | | 0.0008 |
| SZ Avain_Lam | **0.0095** | 0.0031 | 0.0044 | |
| | SZ_in-vivo-H7N9 | SZ_in-vitro-H7N9 | SZ Avain_Lam | SZ Human in vitro_Lam |
| SZ_in-vivo-H7N9 | | 0.0020 | 0.0020 | 0.0019 |
| SZ_in-vitro-H7N9 | **0.0251** | | 0.0006 | 0.0008 |
| SZ Avain_Lam | **0.0250** | 0.0037 | | 0.0009 |
| SZ Human in vitro_Lam | **0.0269** | 0.0097 | 0.0105 | |
| | SZ_in-vivo-H7N9 | SZ_in-vitro-H7N9 | SZ Avain_Lam | SZ Human in vitro_Lam |
| SZ_in-vivo-H7N9 | | 0.0028 | 0.0027 | 0.0027 |
| SZ_in-vitro-H7N9 | **0.0247** | | 0.0008 | 0.0008 |
| SZ Avain_Lam | **0.0252** | 0.0050 | | 0.0007 |
| SZ Human in vitro_Lam | **0.0256** | 0.0062 | 0.0074 | |
| | SZ_in-vivo-H7N9 | SZ_in-vitro-H7N9 | SZ Avain_Lam | SZ Human in vitro_Lam |
| SZ_in-vivo-H7N9 | | 0.0028 | 0.0027 | 0.0027 |
| SZ_in-vitro-H7N9 | **0.0247** | | 0.0008 | 0.0008 |
| SZ Avain_Lam | **0.0252** | 0.0050 | | 0.0007 |
| SZ Human in vitro_Lam | **0.0256** | 0.0062 | 0.0074 | |
| | SZ_in-vivo-H7N9 | SZ_in-vitro-H7N9 | SZ Avain_Lam | SZ Human in vitro_Lam |
| SZ_in-vivo-H7N9 | | 0.0022 | 0.0022 | 0.0021 |
| SZ_in-vitro-H7N9 | **0.0243** | | 0.0006 | 0.0016 |
| SZ Avain_Lam | **0.0249** | 0.0055 | | 0.0017 |
| SZ Human in vitro_Lam | **0.0272** | 0.0179 | 0.0190 | |
| | SZ_in-vivo-H7N9 | SZ_in-vitro-H7N9 | SZ_in-vitro-H7N9_Lam | SZ Avain_Lam |
| SZ_in-vivo-H7N9 | | 0.0032 | 0.0030 | 0.0031 |
| SZ_in-vitro-H7N9 | **0.0309** | | 0.0015 | 0.0017 |
| SZ Human in vitro_Lam | **0.0302** | 0.0148 | | 0.0019 |
| SZ Avain_Lam | **0.0326** | 0.0173 | 0.0193 | |
| | SZ_in-vivo-H7N9 | SZ_in-vitro-H7N9 | SZ Avain_Lam | SZ Human in vitro_Lam |
| SZ_in-vivo-H7N9 | | 0.0022 | 0.0023 | 0.0018 |
| SZ_in-vitro-H7N9 | **0.0105** | | 0.0022 | 0.0019 |
| SZ Avain_Lam | **0.0129** | 0.0099 | | 0.0021 |
| SZ Human in vitro_Lam | **0.0108** | 0.0092 | 0.0113 | |
| | SZ_in-vivo-H7N9 | SZ_in-vitro-H7N9 | SZ Human in vitro_Lam | SZ Avain_Lam |
| SZ_in-vivo-H7N9 | | 0.0040 | 0.0039 | 0.0039 |
| SZ_in-vitro-H7N9 | **0.0257** | | 0.0013 | 0.0007 |
| SZ Human in vitro_Lam | **0.0258** | 0.0061 | | 0.0014 |
| SZ Avain_Lam | **0.0260** | 0.0029 | 0.0068 | |

Within group

0.0312

0.0080

0.0022

Between group*

0.0326

0.0108

0.0029
