## Supplementary figure 4-6 for "Sequencing of clinical samples reveals that adaptation keeps establishing during H7N9 virus infection in humans"

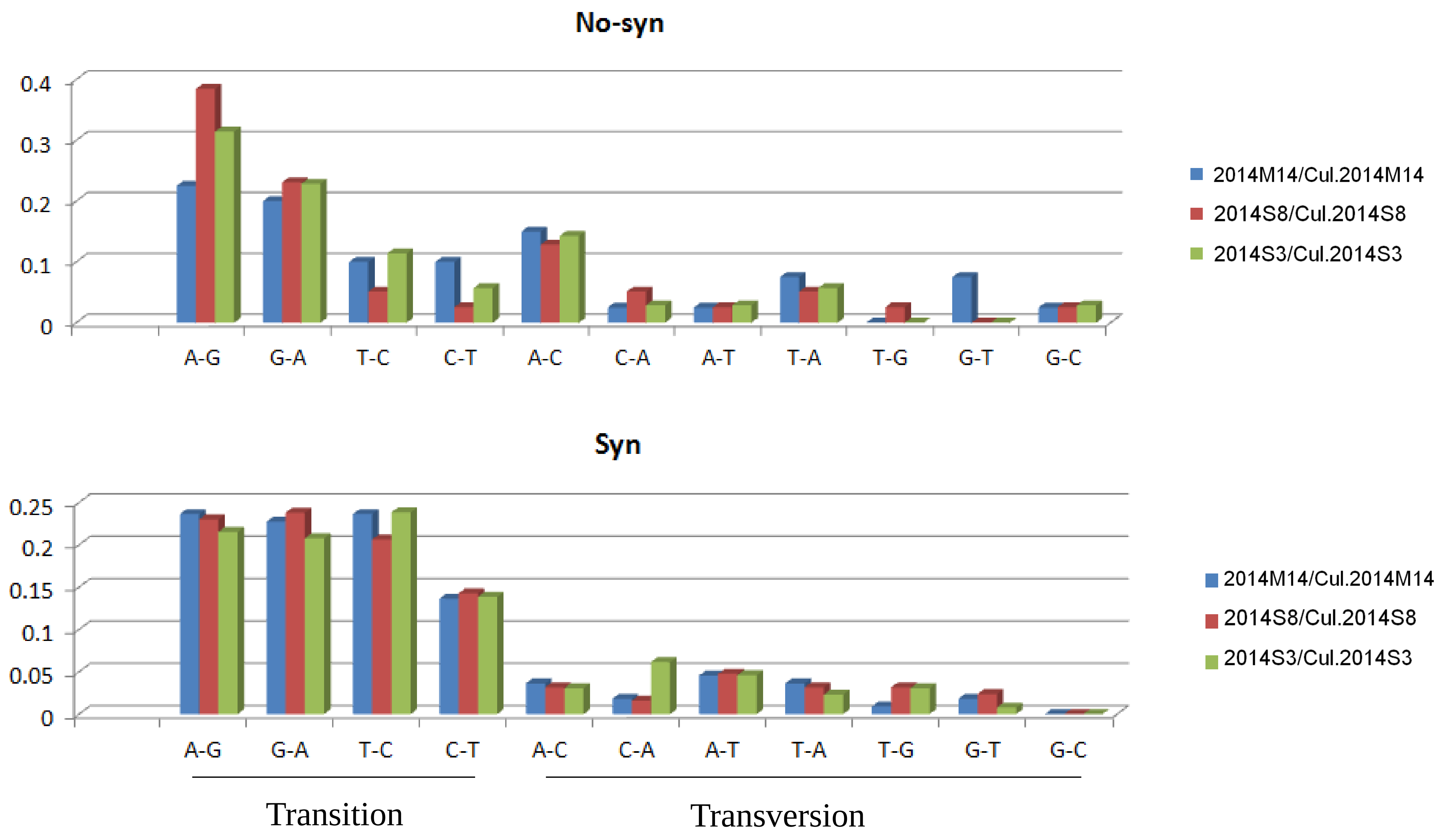

Supplementary figure 5

***In-vivo* human H7N9 VS *in vitro* human H7N9**

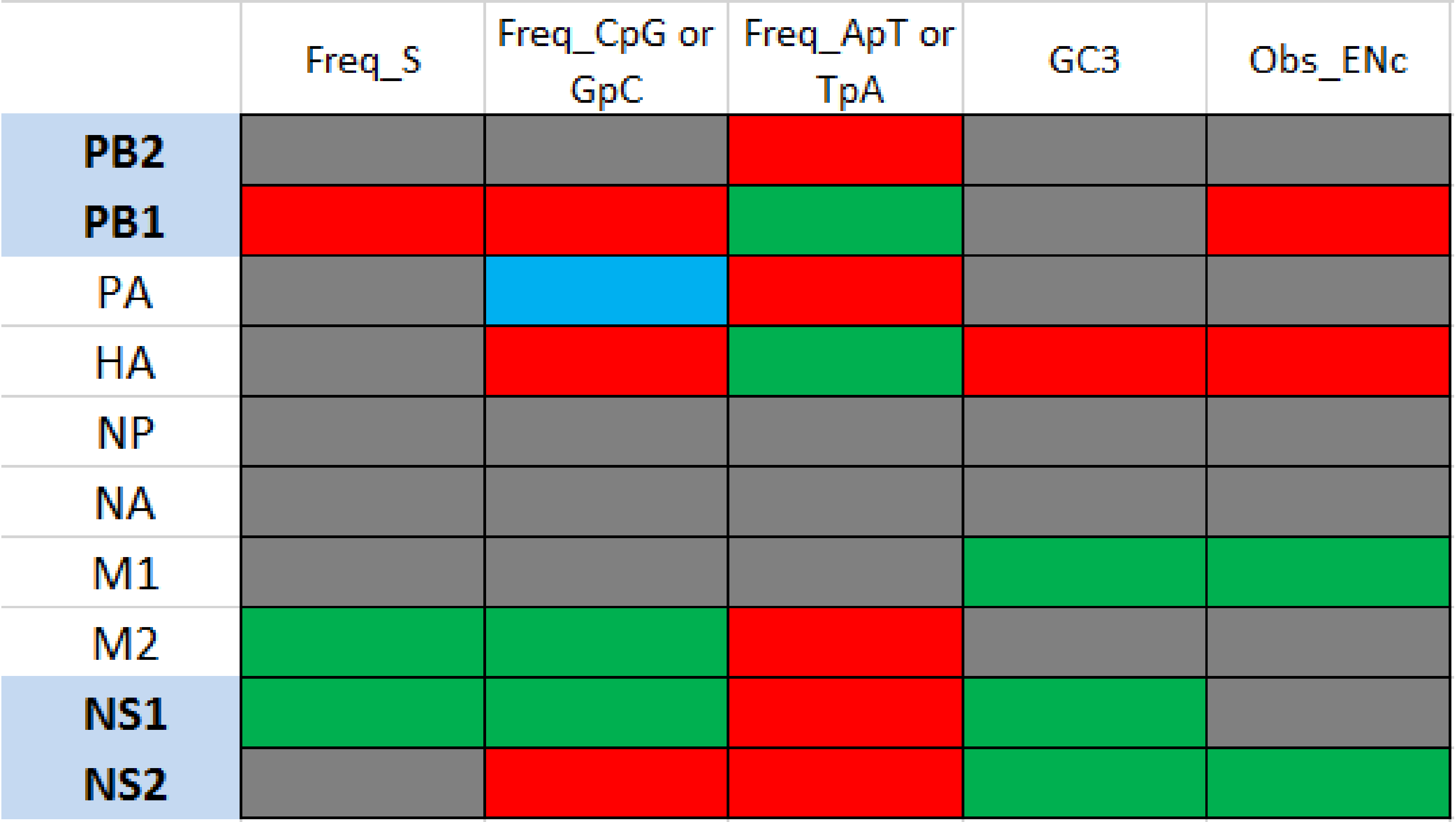

Supplementary figure 6

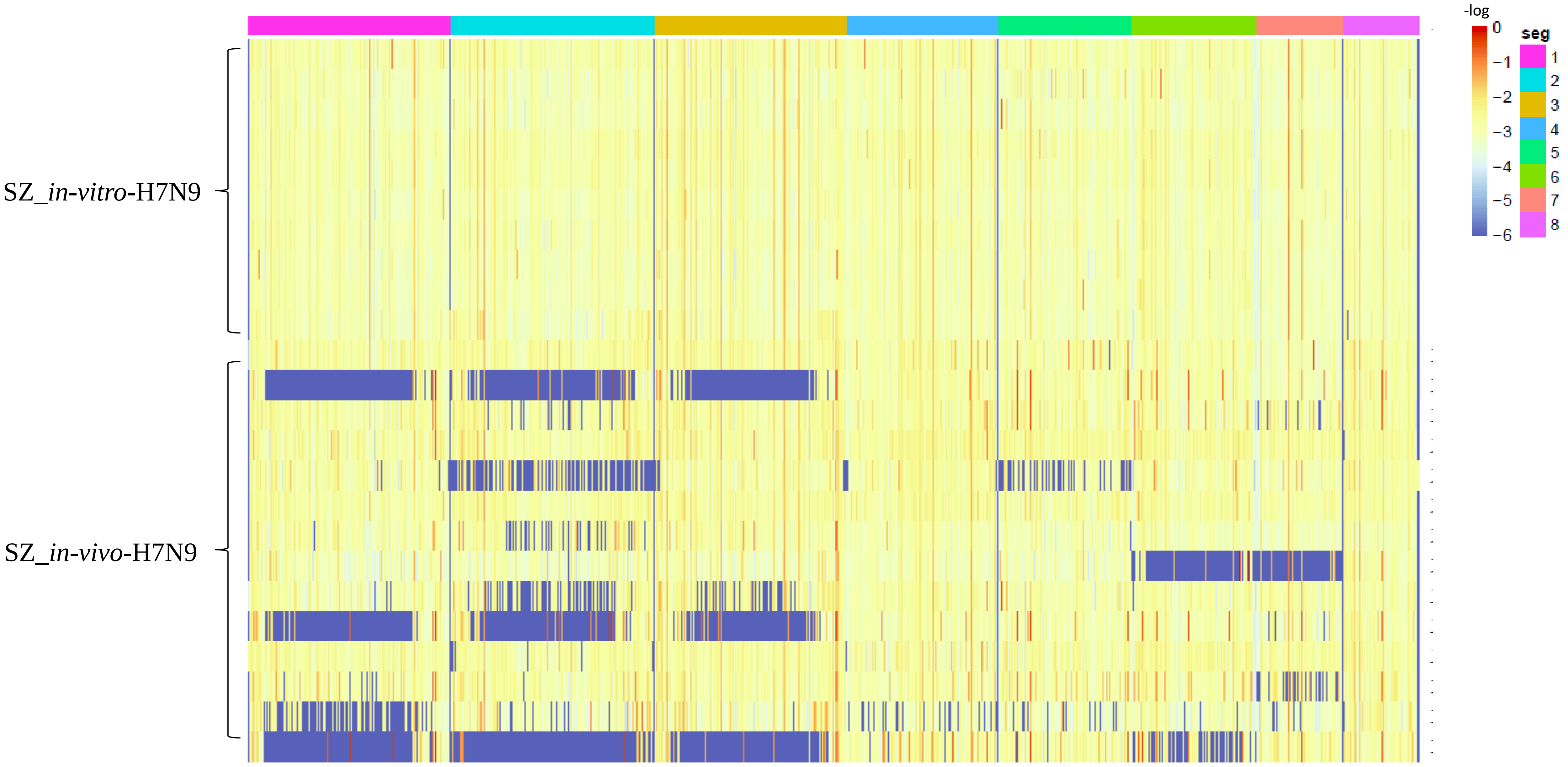
