## Supplementary table 3 for "Sequencing of clinical samples reveals that adaptation keeps establishing during H7N9 virus infection in humans"

Supplemental table 3. Identity of consensus sequence Human H7N9 between *in vivo* samples and from egg cultured ones.

|  | Pairwise length (bp) | 2014M14/Cul.2014M14 | 2014S8/Cul.2014S8 | 2014S3/Cul.2014S3 | 2014S4/Cul.2014S4 |
| --- | --- | --- | --- | --- | --- |
| Seg1 | 2240 | 1.000 | 0.998 | NA | 1.000 |
| Seg2 | 2260 | 0.999 | 0.999 | NA | 1.000 |
| Seg3 | 2145 | 0.999 | 0.997 | NA | 0.999 |
| Seg4 | 1668 | 0.986 | 0.983 | 0.983 | 1.000 |
| Seg5 | 1488 | 0.958 (1407 bp) | 0.960 | 0.952 | 1.000 |
| Seg6 | 1390 | 0.986 (737 bp) | 0.986 | 0.985 | 1.000 |
| Seg7 | 960 | 0.976 | 0.989 | 0.989 | 0.994 |
| Seg8 | 828 | 0.961 | 0.961 | 0.960 | 1.000 |
