## Supplementary table 4 for "Sequencing of clinical samples reveals that adaptation keeps establishing during H7N9 virus infection in humans"

Supplementary Table 4. Synonymous and non-synonymous mutation statistics of 4 pair H7N9 viruses in phase I

|  | 2014S4/Cul.2014S4 | | | 2014M14/Cul.2014M14 | | | 2014S8/Cul.2014S8 | | | 2014S3/Cul.2014S3 | | |
| --- | --- | --- | --- | --- | --- | --- | --- | --- | --- | --- | --- | --- |
|  | No-syn | Syn | dN/dS | No-syn | Syn | dN/dS | No-syn | Syn | dN/dS | No-syn | Syn | dN/dS |
| PB2 | 0 | 0 | *0.00* | 0 | 0 | *0.00* | 0 | 4 | 0.00 | na | na | na |
| PB1 | 0 | 0 | *0.00* | 0 | 2 | 0.00 | 0 | 2 | 0.00 | na | na | na |
| PA | 0 | 1 | 0.00 | 0 | 1 | *0.00* | 3 | 3 | 1.00 | na | na | na |
| HA | 0 | 0 | *0.00* | 9 | 13 | 0.69 | 12 | 15 | 0.80 | 11 | 16 | 0.69 |
| NP | 0 | 0 | *0.00* | 8 | **50** | **0.16** | 5 | **54** | **0.09** | 5 | **65** | **0.08** |
| NA | 0 | 0 | *0.00* | 4 | 6 | 0.67 | 9 | 10 | 0.90 | 8 | 12 | 0.67 |
| M1 | 2 | 2 | 1.00 | 4 | 10 | 0.40 | 2 | 6 | 0.33 | 2 | 6 | 0.33 |
| M2 | 1 | 1 | 1.00 | 9 | 2 | **4.50** | 2 | 1 | **2.00** | 2 | 1 | **2.00** |
| NS1 | 0 | 0 | *0.00* | 7 | **19** | **0.37** | 7 | **19** | **0.37** | 8 | **19** | **0.42** |
| NEP | 0 | 0 | *0.00* | 0 | 9 | **0.00** | 0 | 9 | **0.00** | 1 | 9 | **0.11** |
| Average | 0.3 | 0.4 |  | 4.1 | 11.2 |  | 4.0 | 12.3 |  | 5.3 | 18.3 |  |
| Total | 3 | 4 | 0.75 | 41 | 112 | 0.37 | 40 | 123 | 0.33 | 37 | 128 | 0.29 |
