## Supplementary table 5 for "Sequencing of clinical samples reveals that adaptation keeps establishing during H7N9 virus infection in humans"

Supplementary table 5. Codon-based test of neutrality for analysis of 4 paired H7N9 viruses in phase I.

|  | PB2** | PB1 | PA | HA1 | HA2 | NP | NA | M1 | M2 | NS1 | NS2 |
| --- | --- | --- | --- | --- | --- | --- | --- | --- | --- | --- | --- |
| WHY-G1/ SZCDC-17 | 0;  1.0000 | 0;  1.0000 | -1.0011; 0.3188 | 0;  1.0000 | 0;  1.0000 | 0;  1.0000 | 0;  1.0000 | -0.9365; 0.3509 | 1.4210; 0.1579 | 0;  1.0000 | 0;  1.0000 |
| ZXY-Y1/SZCDC-4 | 0;  1.0000 | -1.4171; 0.1591 | -1.0011; 0.3188 | -1.2292; 0.2214 | -2.7167; 0.0076 | -7.4512;  0 | -1.9208; 0.0571 | -2.7944; 0.0061 | -1.4507; 0.1495 | -3.9438; 0.0001 | -3.2115; 0.0017 |
| ZYQ-A2/ SZCDC-16 | NA | NA | NA | -1.6572; 0.1001 | -2.8063; 0.0058 | -8.7023;  0 | -2.7383; 0.0071 | -2.1979; 0.0299 | 1.7446; 0.0836 | -3.8598; 0.0002 | -3.1102; 0.0023 |
| ZZX/SZCDC-2 | -2.0079; 0.0469 | -1.4171; 0.1591 | -0.9171; 0.3609 | -1.3960; 0.1653 | -2.6982; 0.0080 | -7.8797;  0 | -2.2497; 0.0263 | -2.1979; 0.0299 | 1.7446; 0.0836 | -3.9438; 0.0001 | -3.2115; 0.0017 |

*The probability of rejecting the null hypothesis of strict-neutrality (dN = dS). Values of P less than 0.05 are considered significant at the 5% level and are highlighted in red. The test statistic (dN - dS) is shown above the diagonal. dS and dN are the numbers of synonymous and nonsynonymous substitutions per site, respectively. The variance of the difference was computed using the analytical method. Analyses were conducted using the Nei-Gojobori method [1]. All analysis involved 2 paired nucleotide sequences. All ambiguous positions were removed for each sequence pair. The above analyses were conducted in MEGA6 [2].

** “0; 1.0000” the front values is P value, the other value (1.0000) is the test statistic (dN - dS) value.

1. Nei M. and Gojobori T. (1986). Simple methods for estimating the numbers of synonymous and nonsynonymous nucleotide substitutions. Molecular Biology and Evolution 3:418-426.

2. Tamura K., Stecher G., Peterson D., Filipski A., and Kumar S. (2013). MEGA6: Molecular Evolutionary Genetics Analysis version 6.0. Molecular Biology and Evolution30: 2725-2729.
