## Supplementary table 8 for "Sequencing of clinical samples reveals that adaptation keeps establishing during H7N9 virus infection in humans"

**Supplementary table 8** Functional residues list of H7N9 viral proteins identified in this study.

| Protein | AA site | AAs |  |  |  |
| --- | --- | --- | --- | --- | --- |
| PB2 | 570 | M | 570M |  |  |
| PB2 | 627a | K | 627aK |  |  |
| PB2 | 627c | V | 627cV |  |  |
| PB2 | 701b | D | 701bD |  |  |
| PB2 | 701c | E | 701cE |  |  |
| PB1 | 149 | I | 149I |  |  |
| PB1 | 552 | V | 552V |  |  |
| PA | 57 | Q | 57Q |  | (1) |
| PA | 237 | K | 237K |  |  |
| PA | 262 | K | 262K |  |  |
| PA | 444 | D | 444D |  |  |
| HA | 96(88) | V | 96V |  | **Esterase subdomain** |
| HA | 122(114) | K | 122K | Around 130 loop | receptor binding sub-domain |
| HA | 186(**177**) | I | **186I** |  | receptor binding sub-domain |
| HA | 195(**186**) | A | **195A** | 190-195 hekix | receptor binding sub-domain |
| HA | 282(**273**) | I | 282I |  | **Esterase subdomain** |
| HA | 285(**276**) | D | 285D |  | **Fusion** |
| HA | 300(**291**) | K | 300K |  | **Fusion** |
| NA | 78 | K | 78K |  |  |
| NA | 178 | A | 178A |  |  |
| NA | 320 | S | 320S |  |  |
| NA | 345 | I | 345I |  |  |
| NA | 396 | T | 396T |  |  |
| NA | 431 | G | 431G |  |  |
| NP | 239 | V | 239V |  |  |
| **NP** | **375** | **D** | **375D** |  | (1) |
| NP | 428 | V | 428V |  |  |
| NP | 452 | K | 452K |  |  |
| M1 | 232 | N | 232N |  |  |
| M1 | 248 | M | 248M |  |  |
| M2 | 18 | K | 18K |  |  |
| **M2** | **24** | **E** | **24E** |  |  |
| M2 | 89 | S | 89S |  |  |
| **NS1** | **27** | **M** | **27M** |  |  |
| NS1 | 145 | V | 145V |  |  |
| NEP | 92 | S | 92S |  |  |

Note:

1. AA site in black font is shift AAs in 4 paired culture-uncultured samples and validated by population frequency statistic test, showing differences from natural avian hosts and egg cultured human H7N9 viruses.

2. AA site in bold black font is shift AAs in 4 paired culture-uncultured samples and validated by population frequency statistic test (show differences from natural avian hosts) but show no significant difference from human H7N9 egg culutred ones deposit in genbank.

3. AA site in blue font is AAs showing population frequncy bias between Shenzhen Human H7N9 and egg cultured Shenzhen Human H7N9 viruses, and they all were validated by population frequency statistic test showing differences from natural avian hosts and egg cultured human H7N9 viruses.

1.	**Manz B, Schwemmle M, Brunotte L.** 2013. Adaptation of avian influenza A virus polymerase in mammals to overcome the host species barrier. Journal of virology **87:**7200-7209.
