## Supplementary table 9 for "Sequencing of clinical samples reveals that adaptation keeps establishing during H7N9 virus infection in humans"

**Supplementary table 9.** Correlation analysis among different viral genomic segments of *in vivo* and *in vitro* H7N9 viruses. Pearson correlation coefficient were calculated using average entropy of each genomic segment of every sample from *in vivo* or *in vitro*.

*In vivo*

| Human | PB2 | PB1 | PA | HA | NP | NA | M1/M2 | NS1/NS2 |
| --- | --- | --- | --- | --- | --- | --- | --- | --- |
| PB2 | 1.0000 |  |  |  |  |  |  |  |
| PB1 | 0.7554* | 1.0000 |  |  |  |  |  |  |
| PA | 0.6427 | 0.8175 | 1.0000 |  |  |  |  |  |
| HA | 0.3935 | 0.4854 | 0.5672 | 1.0000 |  |  |  |  |
| NP | 0.5782 | 0.7426 | 0.7465 | 0.7276 | 1.0000 |  |  |  |
| NA | 0.2917 | 0.3121 | 0.3929 | 0.3568 | 0.3981 | 1.0000 |  |  |
| M1/M2 | 0.4744 | 0.5563 | 0.5609 | 0.6164 | **0.7865** | 0.5043 | 1.0000 |  |
| NS1/NS2 | 0.4294 | 0.5183 | 0.5038 | 0.6505 | **0.8704** | 0.4641 | 0.6565 | 1.0000 |

*In vitro*

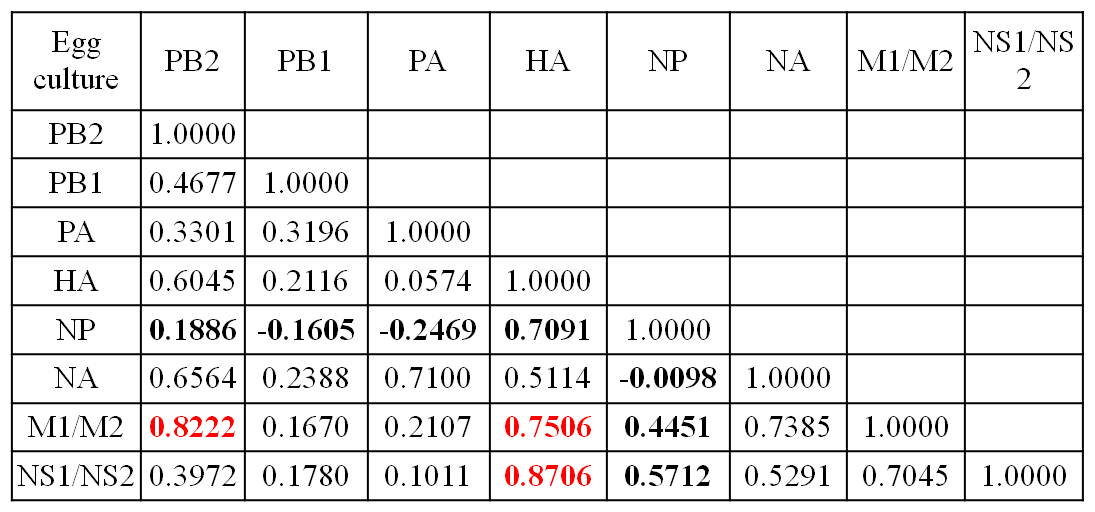

* Pearson correlation coefficients were calculated using average entropy of each genomic segment of every sample from *in vivo* or *in vitro*.
