## Supplementary table 11 for "Sequencing of clinical samples reveals that adaptation keeps establishing during H7N9 virus infection in humans"

Supplemental table 11. Primers for H7N9 genome amplification in this study

| **Primers** | **Sequences (5'-3')** |
| --- | --- |
| uniR_RT | AGTAGAAACAAGG |
| uniF_RT | AGCGAAAGCAGG |
| PB2-F | GATGTCACAGTCTCGCACYC |
| PB2-R | CATCCGAATTCTTTTGGTCGCT |
| PB1-F | GGATGTCAATCCGACTTTRCTYT |
| PB1-R | GTCTGAGCTCTTCAATGGTGGA |
| PA-F | AGACTTTGTGCGACARTGCTTCAAT |
| PA-R | ATCTTAGTGCATGTGTGAGGAAGG |
| H7-F | GAACACTCAAATCCTGGTATTCGC |
| H7-R | GCACCGCATGTTTCCATTCTTTA |
| NP-F | CGWCTCAAGGCACCAAACGA |
| NP-R | TGTCATACTCCTCTGCATTGTCTC |
| N9-F | TCCAAATCAGAAGATTCTATGCACT |
| N9-R | AGGAAGTACTCTATTTTAGCCCCA |
| MP-F | TGAGTCTTCTAACCGAGGTCGAAA |
| MP-R | AAATGACYAKCGTCAACATCCAC |
| NS-F | GCAGGGTGACAAAAACATAATGGA |
| NS-R | ACGAGAAAGTTCTTATCTCTTGCT |
